# IRF7 deficiency increases disease severity independently of TLR7 recognition in Influenza A infection in mice

**DOI:** 10.64898/2026.05.23.727394

**Authors:** Ana Karina Nisperuza Vidal, Chenxiao Wang, Michaela J Allen, Xiaofeng Ding, Jefferson Evangelista, Yilin Chen, Mst Shamima Khatun, Amrita Kumar, Calder Ellsworth, Mohammad Islamuddin, Robert Blair, Suryaprakash Sambhara, Jay K Kolls, Derek A Pociask, Xuebin Qin

## Abstract

Influenza A virus (IAV) remains a major cause of respiratory morbidity and mortality, yet the role of Toll-like receptor 7 (TLR7), an RNA sensor, and its downstream signaling events, such as interferon regulatory factor 7 (IRF7), in IAV infection remain unclear. To address this question, we used single-cell RNA sequencing, genetic mouse models, and immunological analysis. Single -cell transcriptomic profiling of the infected lungs revealed robust upregulation of *Tlr7* and genes associated with interferon pathways in dendritic cells and B cells, alongside widespread induction of *Irf7* across immune and non-immune compartments. *Tlr7*-deficient mice exhibited normal viral control, lung pathology, and survival following IAV challenge. In contrast, *Irf7* deficiency resulted in significantly increased disease severity, impaired early interferon responses, exacerbated bronchial epithelial hyperplasia, and defective early humoral priming. In assessing adaptive immunity, both *Irf7*-deficient and*Tlr7*-deficient mice had reduced antihemagglutinin antibody production. Mechanistically, IRF7 protein expression and downstream signaling were largely preserved in TLR7-deficient mice, indicating that IRF7 activation during IAV infection occurs independently of TLR7. Collectively, these findings identify IRF7 as a non-redundant determinant of innate immunity and disease outcomes during IAV infection, while positioning TLR7 as a modulatory factor primarily influencing adaptive immune maturation. Our study refines current models of antiviral sensing by uncoupling receptor induction from functional necessity and highlights IRF7 as a critical downstream regulator dictating host defense against acute influenza A infection.

**Importance:** Influenza A virus is a respiratory pathogen that remains a major threat to global health as a seasonal disease and a source of periodic pandemics. The outcomes of the infection can range from mild illness to severe pneumonia and death, particularly in vulnerable populations, yet the reasons why some individuals develop more severe disease are not fully understood. Early immune defenses in the lungs are critical for controlling the virus, but they can also contribute to harmful inflammation if not properly regulated. In particular, key sensors that detect viral genetic material and the signaling pathways that activate antiviral responses play an essential role in shaping these outcomes. The significance of our study lies in defining how these early immune mechanisms influence the course of influenza A infection, providing insight that may guide the development of improved therapies for influenza and related respiratory viruses.

## Introduction

The RNA viruses SARS-CoV-2 (CoV2) and influenza A (IAV) are respiratory viruses that continue to pose a global threat^1,2^. CoV2, a non-segmented, single-stranded, positive sense RNA virus, infects host cells via its spike (S) protein binding to the angiotensin-converting enzyme 2 (ACE2)^3–5^. In contrast, IAV is a segmented, negative sense, single-stranded RNA virus that infects host cells via its hemagglutinin (HA) binding to sialic acid receptors^3,6^. Although both viruses infect the respiratory epithelium, further investigation is needed to understand the similarities and differences in their pathogenesis and host immune responses, which will inform the development of more effective interventions for the diseases^3^.

Toll-like receptors (TLRs) represent a class of evolutionarily conserved proteins pivotal in the innate immune system’s arsenal against pathogens, including CoV2 and IAV viruses^7,8^. By bridging innate and adaptive immunity, TLRs elicit a comprehensive immune response, ensuring a rapid and efficient reaction to microbial invasions^9^. Among the 10 TLRs in humans, the endosomal TLRs 3, 7, and 8 are critical to sensing RNA viruses^10^. TLR3 and TLR7 protein structures and functions are conserved in mice and humans^11–13^, and they play crucial roles in recognizing viral nucleic acids and triggering innate immune responses, particularly in the production of type I interferons (IFN). Emerging clinical evidence indicates that the mutations of TLR7, encoded by an X-chromosome locus^14^, contribute to the development of severe COVID-19 via reduced interferon IFN response, mainly in plasmacytoid dendritic cells (pDCs)^15–19^. The rare loss-of function mutations of the TLR7 gene attribute to approximately 1-2% of severe COVID-19 cases in males under the age of 60 without medical history^14–19^. Individuals carrying the mutation of TLR7 downstream signaling molecule IRF7^20^, are also prone to viral infections of the respiratory tract^21^. Moreover, autoantibodies against type I interferon correlate with cellular immune alterations in severe COVID-19 in patients, which indicates the reduced IFNs’ function may contribute to the severity of CoV2 infection^22^. Consistently, we documented that the deficiencies of either *Tlr7* or *Irf7* in mice increase the severity of COVID-19 through the reduced interferon production resulting in a substantial reduction in the production of antibodies against CoV2^23^. These findings highlight the importance of TLR7 and the downstream molecules such as IRF7 in innate and adaptive immunity in COVID-19. In contrast, TLR7’s role in IAV infection is less appreciated as compared to COVID-19.

Previous studies emphasize the importance of TLR7 in mediating effective immune responses to IAV infection, particularly in promoting adaptive immunity and enhancing antiviral defenses^24^. Other research suggests that TLR7 is not critical in fighting IAV infection, with evidence showing that its absence has minimal impact on viral clearance or immune modulation^25^. A TLR7 antagonist restricts IFN-dependent immunopathology in a mouse model of severe IAV infection^26^. TLR7 promotes acute inflammation and lung dysfunction in mice infected with IAV but prevents late airway hyperresponsiveness^27^. These findings suggest that the role of TLR7 in innate immunity against IAV infection differs from its function in CoV2 infection, underscoring the need for further research to clarify the conditions under which TLR7 significantly contributes to immune defense mechanisms. Conversely, studies find that the activation of TLR7 and IRF7 promotes a broad immune response against heterologous strains of IAV and CoV2^28^. This activation is important for the induction of IAV hemagglutinin (HA)- specific Abs in response to the 2009 pandemic split vaccine in mice^25^, and the amplification of type I and III IFNs is required for protection against primary infection by IAV in humans^29^. The consensus is that TLR7 is dispensable for disease recovery in the acute stage of disease^25^. Nevertheless, the extent to which TLR7 and its downstream signaling contribute to the innate and adaptive immune responses to IAV remain poorly defined. To address this question, we mined our previously published single-cell RNA sequence (scRNA-seq) data^30^ obtained from the lungs of the mice either infected with or without the IAV PR8 (A/Puerto Rico/8/1934H1N1) infection for exploring the transcriptional changes of TLRs and related signaling pathways at 4 and 6 days post-infection (DPI). We also utilized the TLR7 and IRF7-deficient mice to explore the role of TLR7 and IRF7 in the immune response to the infection of PR8, a well characterized mouse adapted strain.

## Results

### IAV infection induces up-regulation of *Tlr7* transcripts

To define baseline expression patterns, we reanalyzed the Tabula Muris Consortium single cell RNAseq dataset from lungs of four 10–15 week old male and four virgin female C57BL/6JN mice^31^. *Tlr7* expression was mainly expressed in classical, non-classical monocyte subsets, and alveolar macrophages (Fig. 1A and Supplementary. Table 1A). In contrast, *Tlr3* expression was more broadly distributed across multiple cell types, including, alveolar macrophages, myeloid cells, endothelial and stromal compartments (Fig. 1B and Supplementary. Table 1B). These results indicate a more cell type–specific pattern for *Tlr7* versus a wider but sparse expression of *Tlr3* under steady state conditions.

**Figure 1.**
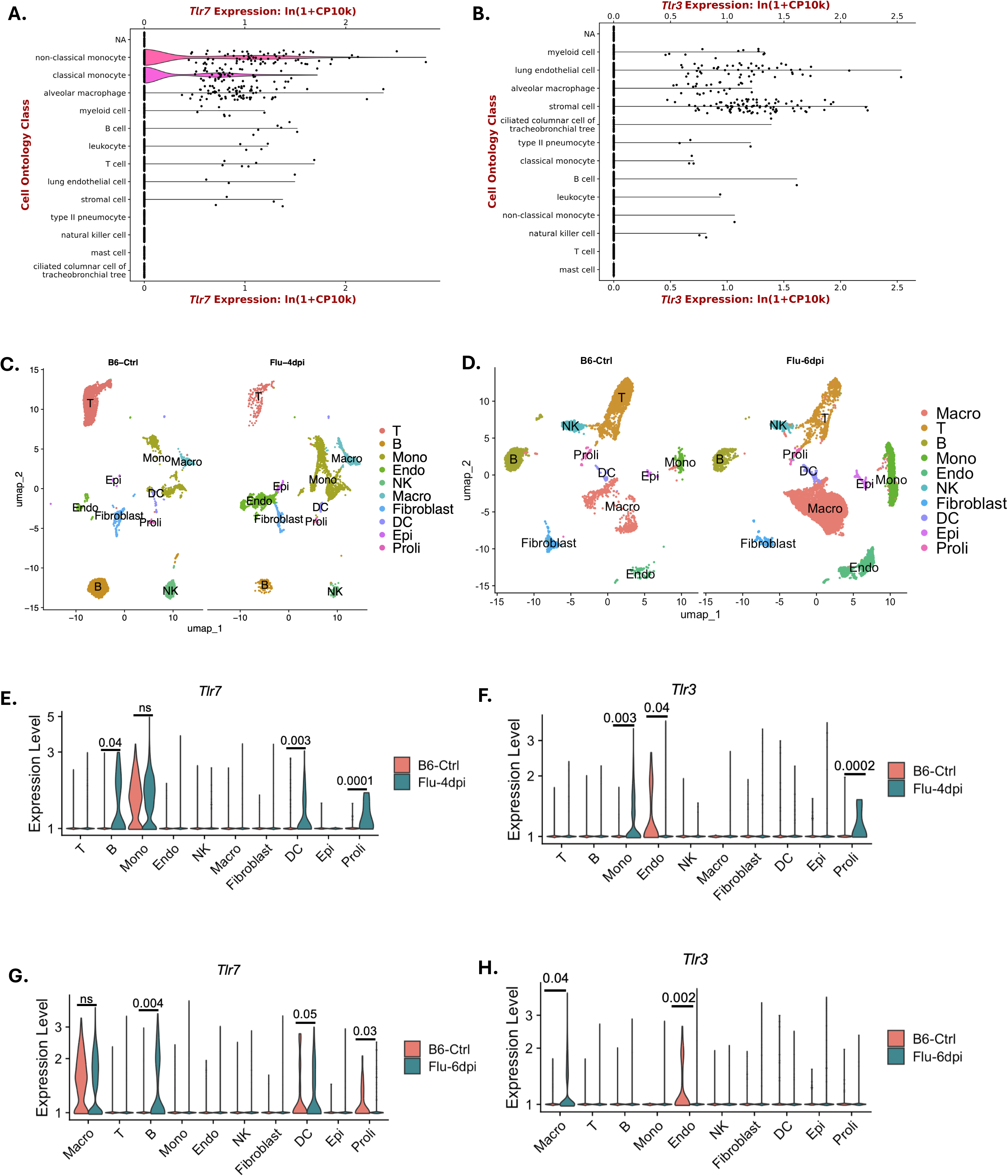
Upregulation of Tlr7 during IAV infection. **A–B)** Expression of *Tlr7* (A) and *Tlr3* (B) across lung cell populations from the droplet-based single-cell RNA-seq dataset of the Tabula Muris Consortium^31^. **C-D)**UMAP visualization of major lung cell populations from two 12-week-old B6 control and two 12-week-old noninfected B6 mice IAV infected mice at 4 dpi (C) and 6 dpi (D), based on single-cell RNA-seq analysis. **E-F)** Expression of *Tlr7* (C) and *Tlr3* (D) across lung cell populations in control and IAV infected mice at 4 dpi. **G-H)** Expression of *Tlr7* (G) and *Tlr3* (H) across lung cell populations in control and influenza-infected mice at 6 dpi. Expression levels are shown as normalized single cell expression values. Statistical significance between groups is indicated above each cell population*; ns, not significant*.

To investigate the expression patterns and transcriptomic changes induced by IAV on *Tlr7*, we mined our previously published scRNA-seq data of mice lung tissue from uninfected and IAV-infected mice^30^. The infection was established using 50 PFU IAV PR8 (A/Puerto Rico/8/1934H1N1) administered intranasally. *Tlr7* and *Tlr3* expression were assessed at the single-cell level by analyzing scRNA-seq of the lungs collected from PR8-infected mice at 4- and 6-day post-infection (DPI)^30^. Using uniform manifold approximation and projection (UMAP) for dimensionality reduction, we identified 10 distinct cellular clusters in both timepoints (Fig. 1C, D). At 4 DPI, *Tlr7* was significantly upregulated in B cells and dendritic cells (DCs) (Fig. 1C), whereas statistically significant differences in *Tlr3* expression were observed in monocytes (Fig. 1D). This pattern of expression was consistent at 6 DPI, with a higher *Tlr7* expression in B and DC cells (Fig. 1G). and a higher *Tlr3* expression in macrophages of PR8-infected lungs as compared to uninfected lungs (Fig. 1H). Collectively, these findings demonstrate that there is a consistent upregulation of *Tlr7* in pulmonary immune cells during IAV infection.

### *Tlr7* deficiency does not increase disease severity in IAV infection

To elucidate the role of *Tlr7* in the development of IAV infection pathology, we utilized male mice, 12-16 weeks (about 3 and a half months) old, *Tlr7*-deficient (*Tlr7*^−/−^) and *Tlr7*-sufficient (*Tlr7*^+/+^). Animals were intranasally inoculated with a lethal dose of 83 PFU of the PR8 influenza A strain. Following infection, we tracked daily changes in body weight (Fig. 2a) and assessed survival outcomes (Fig. 2B). Both Tlr7 and *Tlr7*^−/−^ mice exhibited progressive weight loss up to 8 DPI with a good recovery starting at 9 DPI (Fig. 2A). No differences in survival rates were observed (Fig. 2B). Notably, viral RNA quantity was indistinguishable between genotypes (Supplementary Fig. 1). PR8 infection resulted in pulmonary pathology by 7 DPI, but histopathologic analysis did not reveal any significant differences regarding bronchial damage, interstitial inflammation, edema, or bronchial epithelial hyperplasia between controls and *Tlr7*^−/−^ mice (Fig. 2C-H). These findings suggest that *Tlr7* deficiency does not affect the recovery from IAV infection.

**Figure 2.**
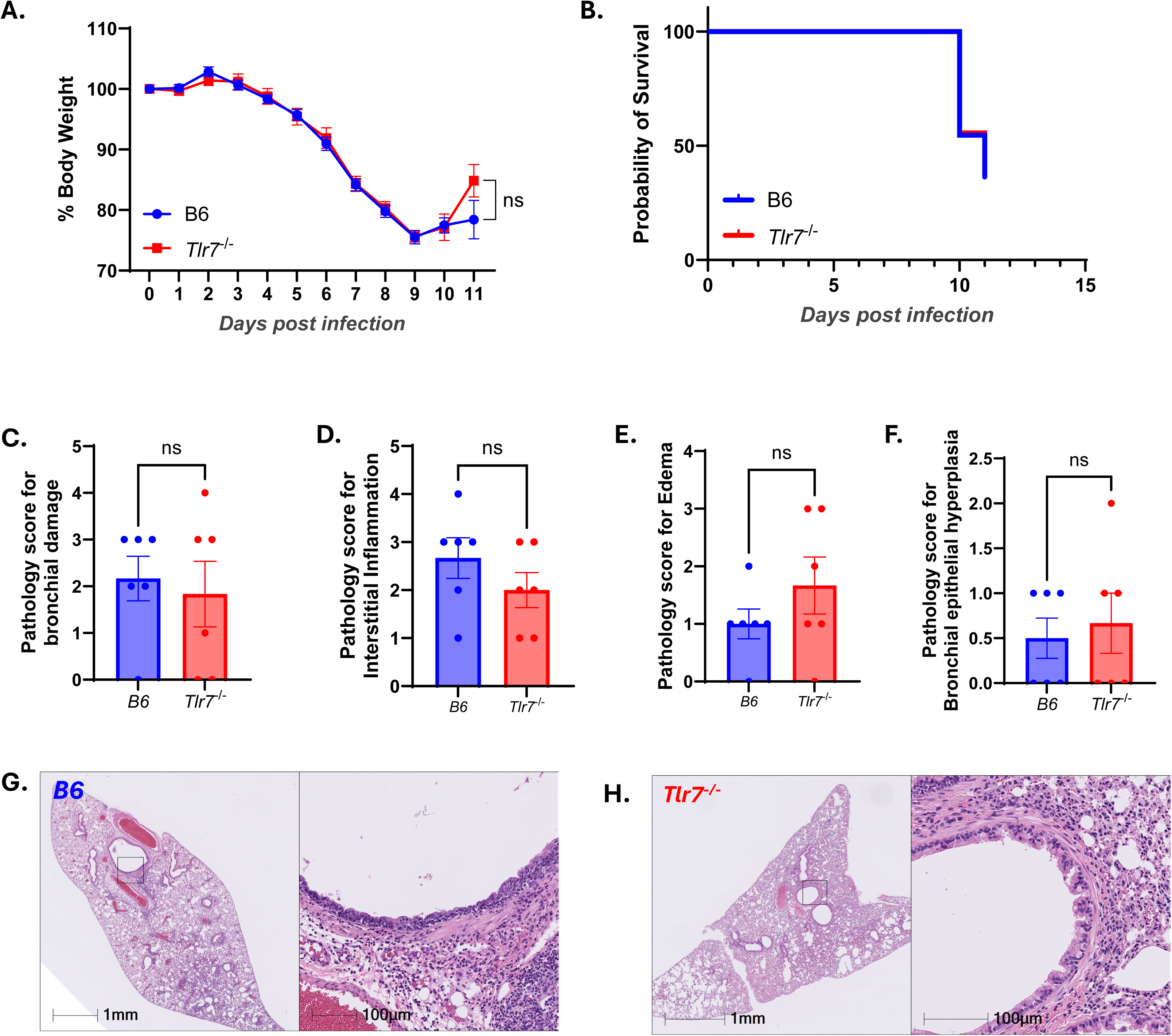
Global *Tlr7* deficiency does not affect mice recovery in IAV infection. **A)** Male *Tlr7^−/−^* mice did not show differences in body weights compared to age-matched male B6 mice PR8 infection. 12-week-old male B6 mice (*n* = 9) and age-matched B6 (*n* = 11) male mice were intranasally inoculated with Influenza PR8 infection (83 PFU). Body weights were monitored daily. Results are shown as mean ± SEM. The body weight difference is analyzed by a mixed-effects model. *p* = 0.5629. **B)** Survival analysis of male *Tlr7^−/−^* mice and age-matched male B6 mice post PR8 infection. 12-week-old male *Tlr7^−/−^* mice (*n* = 9) and age-matched B6 (*n* = 11) male mice were administrated with PR8 infection (83 PFU). The comparison of survival curves is analyzed by the Log-rank (Mantel-Cox) test. *p* = 0.7632. **C – H)** Representative images and quantitative histology analysis of H and E staining of infected B6 and *Tlr7^−/−^* mice. 12-week-old male *Tlr7^−/−^* mice (*n* = 6) and age-matched B6 (*n* = 6) at 9 - 11 DPI were included for analysis. No differences were found in the pathology analysis for bronchial damage, interstitial inflammation, edema and bronchial epithelia hyperplasia between *Tlr7^−/−^* and B6 mice. Pathology comparisons are analyzed by unpaired *t*-test and are presented as mean ± SEM. *P < 0.05, **P < 0.01, and ***P < 0.001. *ns, not significant*.

### *Irf7* expression during IAV infection

The Gene Set Enrichment Analysis (GSEA) of IAV-infected mice showed upregulation of genes related to the Interferon gamma (IFN-γ) and Interferon alpha (IFN-α) responses at 4 and 6 DPI in B cells (Fig. 3 A, B) and dendritic cells (DCs) (Fig. 3. C, D) and monocytes (Fig 3. E, F). These are the same cell populations where *Tlr7* was regulated (Fig. 1. C, E). Additionally, GSEA showed IFN-γ and IFN-α responses upregulated in macrophages and epithelial and endothelial cells (Supplementary Fig. 2). Taking into account the GSEA results, and that the transcription factor downstream of TLR7 is IRF7, and that we previously have demonstrated the importance of the TLR7-IRF7 axis in the context of another RNA respiratory virus, SARS-CoV-2^23^, we further explored the expression levels of IRF7 in our single-cell data. *Irf7* expression dramatically increased in the IAV-infected group both at 4 and 6 DPI (Fig. 4. A, B) in monocytes, macrophages, DCs, B cells, T cells, NK, epithelial, and endothelial cells and fibroblasts. On the contrary, *Irf3* (Fig. 4. C, D) and *Irf8* (Fig. 4. E, F) did not show an increased expression as *Irf7*. These findings suggest that TLR7 regulates IRF7 during IAV infection, influencing IFN-α and IFN-γ pathway activation in immune cells.

**Figure 3.**
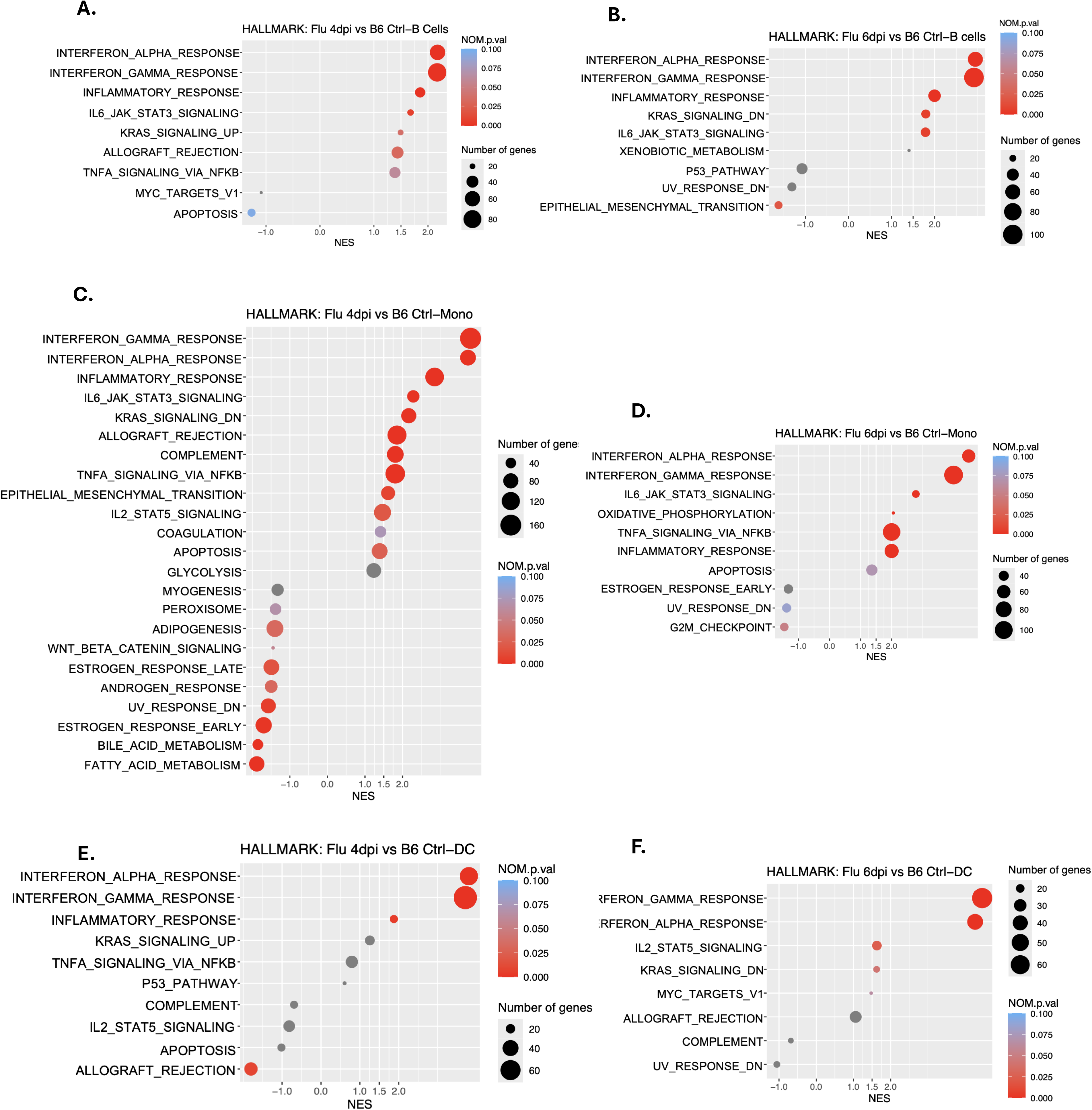
Upregulation of IFN responses in IAV infected mice. **A–B)** Gene set enrichment analysis (GSEA) of Hallmark pathways in B cells from IAV infected versus B6 control mice at 4 dpi (A) and 6 dpi (B). **C–D)** GSEA of Hallmark pathways in monocytes comparing IAV infected and B6 control mice at 4 dpi (C) and 6 dpi (D). **E–F)** GSEA of Hallmark pathways in dendritic cells comparing IAV infected and B6 control mice at 4 dpi (E) and 6 dpi (F). Normalized enrichment scores (NES) are shown on the x-axis. Dot size represents the number of genes contributing to each pathway, and color indicates normalized *p*-value (NOM p.val).

**Figure 4.**
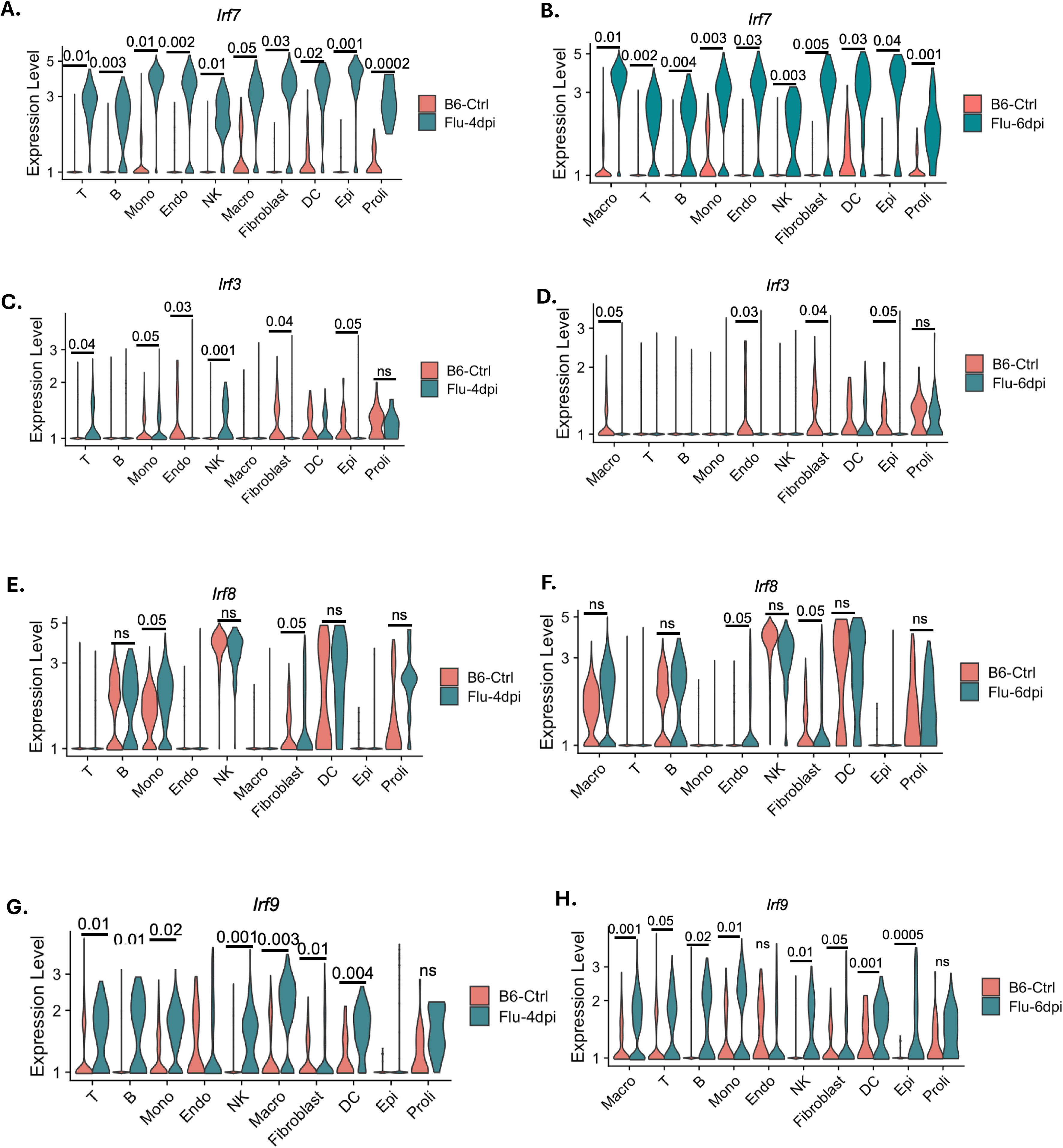
Expression of different *Irf* during IAV infection. **A–B)** Single-cell RNA-seq analysis of *Irf7* expression across indicated lung cell populations in B6 control and influenza-infected mice at 4 dpi (A) and 6 dpi (B). **C–D)** Expression of *Irf3* across lung cell populations at 4 dpi (C) and 6 dpi (D). **E–F)** Expression of *Irf8* across lung cell populations at 4 dpi (E) and 6 dpi (F). **G–H)** Expression of *Irf9* across lung cell populations at 4 dpi (G) and 6 dpi (H). Expression levels are shown as normalized single cell expression values. Statistical significance between groups is indicated above each cell population; *ns, not significant*.

### Protective role of *Irf7* in IAV infection

To further explore the role of Irf7 in IAV infection, we infected *Irf*^−/−^ mice, 12 to 16 weeks old, *Irf7*-deficient (*Irf7*^−/−^) and *Irf7*-sufficient (*Irf7*^+/+^) with a lethal dose of 83 PFU of PR8. Infected Irf7 lost more body weight up to 9 DPI and showed a higher mortality rate than the sufficient *Irf7*^+/+^ controls (Fig. 5A, B). Histological studies demonstrated that *Irf7*^−/−^ mice had significantly higher bronchial epithelial hyperplasia (Fig. 5F, H). However, no differences were found when comparing other histopathologic changes such as bronchial damage, interstitial inflammation, and edema (Fig. 5C – E). No significant differences in viral load were observed between control and *Irf7*⁻/⁻ mice at 2 DPI, as assessed by qPCR and immunofluorescence (Fig. 5 I – L). These results indicate that IRF7 influences disease outcome during IAV infection, affecting survival and epithelial responses without altering viral burden.

**Figure 5.**
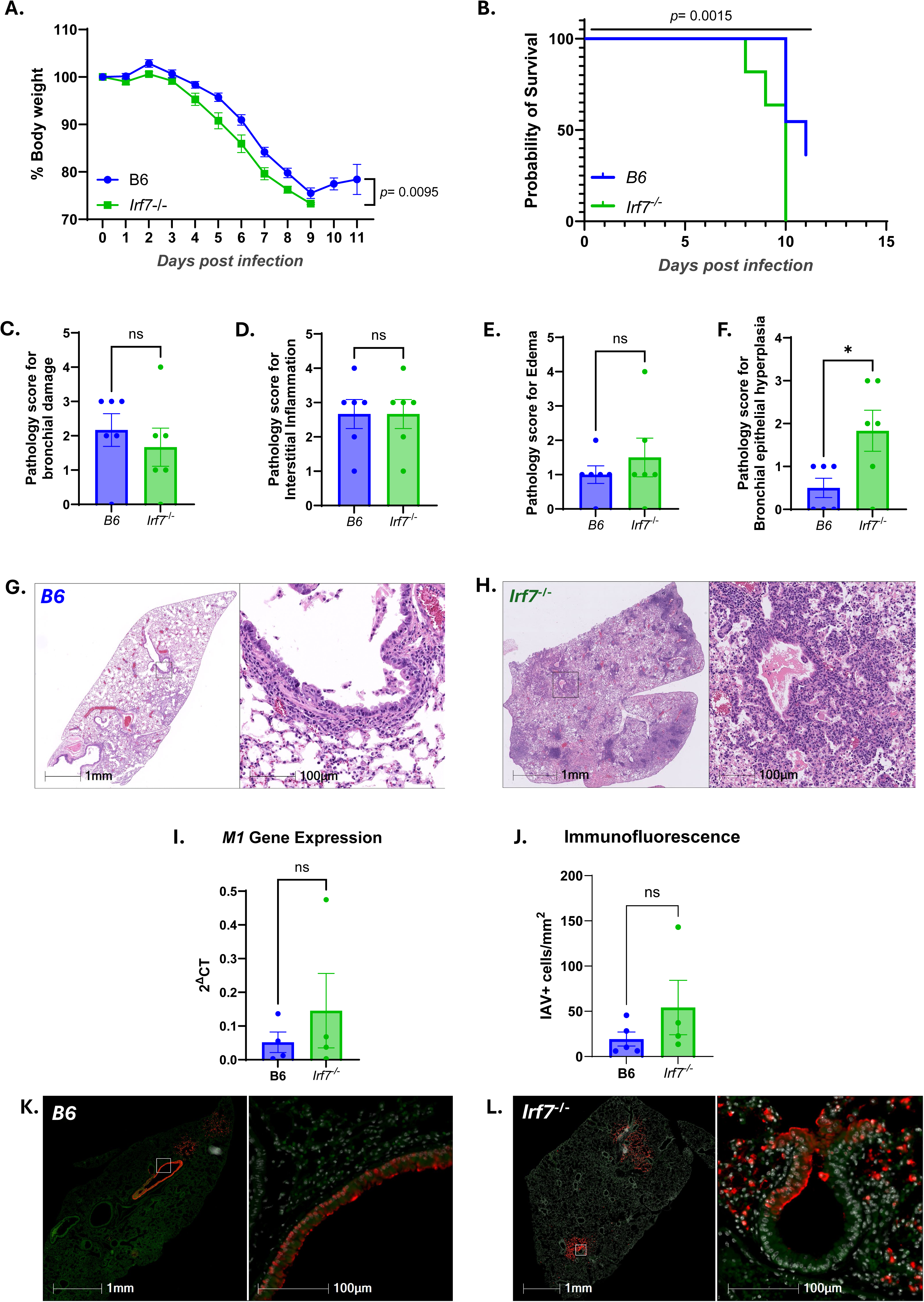
Global Irf7 deficiency increases the severity of IAV infection in mice. Male *Irf7^−/−^*mice lost more body weight compared to age-matched male B6 mice PR8 infection. 12-week-old male B6 mice (*n* = 11) and age-matched B6 (*n* = 11) male mice were intranasally inoculated with Influenza PR8 infection (83 PFU). Body weights were monitored daily. Results are shown as mean ± SEM. The body weight difference is analyzed by a mixed-effects model (*p* =0.0095). **B)** Survival analysis of male *Irf7^−/−^* mice and age-matched male B6 mice post PR8 infection. 12-week-old male *Irf7^−/−^* mice (*n* = 11) and age-matched B6 (*n* = 11) male mice were administrated with PR8 infection (83 PFU). The comparison of survival curves is analyzed by the Log-rank (Mantel-Cox) test (*p* = 0.0015) **C – H)** Representative images and quantitative histology analysis of H and E staining of infected B6 and *Irf7^−/−^* mice. 12-week-old male *Irf7^−/−^*mice (*n* = 11) and age-matched B6 (*n* = 11) at 7 - 11 DPI DPI were included for analysis. Significant difference were found in bronchial epithelia hyperplasia between *Irf7^−/−^* and B6 mice *(p* = 0.0388). **I)** Quantification of influenza A M1 gene expression in lung homogenates of infected B6 and *Irf7^−/−^* mice at 2 DPI by RT-qPCR, shown as 2^-ΔCt. **J)** Quantification of IAV positive cells in lung sections by immunofluorescence, expressed as cells/mm². **K–L)** Representative immunofluorescence images of lung sections from B6 (K) and *Irf7^−/−^* (L) mice showing IAV staining (red). Scale bars, 1 mm (left) and 100 µm (right). Pathology comparisons are analyzed by unpaired *t*-test and are presented as mean ± SEM. *P < 0.05, **P < 0.01, and ***P < 0.001. *ns, not significant*.

### The role of *Tlr7 and Irf7* in adaptative immunity during IAV infection

Knowing that *Tlr7* is dispensable for IAV infection recovery and *Irf7* offers a protective effect against it, we further wanted to evaluate their effect in the adaptive immune response following IAV infection. IgG against the hemagglutinin (HA) protein of IAV was measured by the ELISA method in mice infected with lethal (83 PFU of PR8) and sublethal doses (50 PFU of PR8) of IAV. We found that at 7 DPI after a lethal dose, the infected *Irf7*^−/−^ showed a significantly lower level of anti-HA IgG in respect to the control group, while *Tlr7*^−/−^, also showed a lower level than controls, but the difference did not achieve statistical significance (Fig. 6A). Following infection with a sublethal dose that allowed the mice to survive the acute phase of the disease, no differences were found among the B6 controls, *Tlr7*^−/−^, and *Irf7*^−/−^ groups on day 14 post-infection (Fig. 6B). At 21 DPI *Tlr7*^−/−^ showed lower levels of anti-HA IgG compared to the control group, while *Irf7*^−/−^ did not show any differences (Fig. 6C). These findings suggest a priming effect of *Irf7*^−/−^ on antibody production during the earlier phase of infection, while *Tlr7*^−/−^ acts in both the early and late stages on the adaptive immune response.

**Figure 6.**
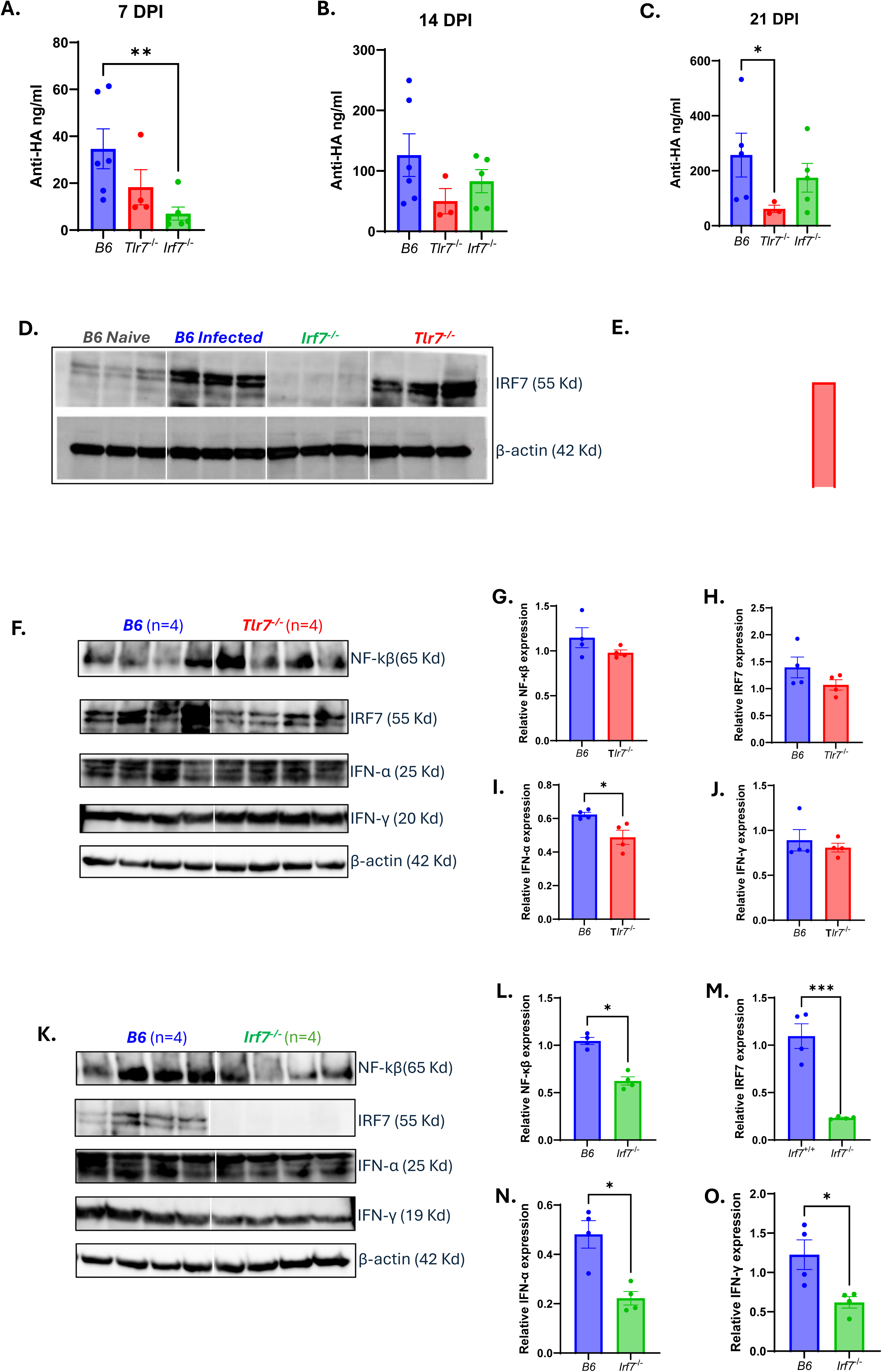
*Tlr7 and Irf7* deficiencies impair adaptative immunity against IAV. **A – C)** Comparison of antibodies against hemagglutinin (Anti-HA, ng/ml) between B6, *Tlr7^−/−^* and *Irf7^−/−^* mice at 7, 14 and 21 DPI. A) *p* = 0.0087 between B6 and *Irf7^−/−^* at 7 DPI. C) *p* = 0.0357 between B6 and *Tlr7^−/−^* at 21 DPI. **D)** Western blot of IRF7 (55 kDa) in lung lysates from B6 naïve, B6 infected, *Irf7*^⁻/⁻^, and *Tlr7*^⁻/⁻^ mice at 7 DPI. β-actin (42 kDa) is shown as a loading control. **E)** Densitometric quantification of IRF7 protein levels normalized to β-actin and expressed relative to B6 naïve controls. **F)** Western blots of NF-κB (65 kDa), IRF7 (55 kDa), IFN-α (25 kDa), and IFN-γ (20 kDa) in lung lysates from B6 and *Tlr7*⁻/⁻ mice at 2 DPI (n = 4 per group). β-actin (42 kDa) is shown as a loading control. **G–J)** Densitometric quantification of NF-κB (G), IRF7 (H), IFN-α (p=0.0286) (I), and IFN-γ (J) protein levels normalized to β-actin and expressed relative to B6 controls. **K)** Western blots of NF-κB (65 kDa), IRF7 (55 kDa), IFN-α (25 kDa), and IFN-γ (20 kDa) in lung lysates from B6 and *Irf7*⁻/⁻ mice at 2 DPI (n = 4 per group). β-actin (42 kDa) is shown as a loading control**. L–O)** Densitometric quantification of NF-κB (p=0.0286) (L), IRF7 (p=0.0049) (M), IFN-α (N) (p=0.0285), and IFN-γ (p=0.0286) (O) protein levels normalized to β-actin and expressed relative to B6 controls. Data are presented as mean ± SEM. Statistical significance was determined by unpaired *t*-test. *P < 0.05, **P < 0.01, and ***P < 0.001. *ns, not significant*.

### Interferon production is reduced in *Irf7*- and *Tlr7*-deficient mice during IAV infection

To assess how *Tlr7* and *Irf7* deficiencies affect IFN production at the protein level during IAV infection, we first measured IRF7 expression by Western blot at 7 - 9 DPI. As expected, IRF7 protein levels were markedly reduced in *Irf7*^−/−^ mice compared to infected B6 controls. In contrast, IRF7 protein expression in *Tlr7*^−/−^ mice was comparable to that observed in infected B6 mice at this time point (Fig. 6D, E). Given that interferon signaling is known to be initiated early during IAV infection, we next evaluated IRF7 and downstream signaling at an earlier stage. At 2 DPI, IRF7 and NF-κB protein levels were similar between infected wild-type and *Tlr7*^−/−^ mice, whereas *Irf7*^−/−^ mice exhibited a significant reduction in IRF7 expression as expected (*p*=0.0049) and a concomitant decrease in NF-κB activation (*p*=0.0286) compared to wild-type controls (Fig. 6F-O). Consistent with these findings, analysis of interferon levels revealed a significant reduction in IFN-α in *Tlr7*^−/−^ mice compared to B6 controls (*p*=0.0286), while IFN-γ levels were not significantly different between these groups. In contrast, *Irf7*^−/−^ mice showed significantly reduced levels of both IFN-α (*p*=0.0285) and IFN-γ (*p*=0.0286) relative to infected B6 mice (Fig. 6F – O). These results indicate that early interferon production during IAV infection is strongly dependent on IRF7, while TLR7 deficiency has a limited impact on IRF7 protein expression and downstream signaling at early and late stages of infection.

## Discussion

In this study, we define a functional dissociation between viral sensing and disease outcome during IAV infection, identifying IRF7 as a critical determinant of innate immunity and disease outcomes during IAV infection, independently of TLR7 signaling. Despite robust induction of *Tlr7* transcripts across multiple immune populations, *Tlr7^−/−^*mice exhibited weight loss, survival, viral RNA levels, and lung histopathology comparable to the wild-type controls, indicating that TLR7 is dispensable for controlling viral burden and acute disease consistent with prior studies demonstrating redundancy of TLR7 in influenza sensing and antiviral control^25,32^. In contrast, *Irf7^−/−^* mice developed increased disease severity with higher body weight loss, mortality, and bronchial epithelial hyperplasia without a corresponding increase in viral load, revealing an independent relation between viral replication and tissue pathology. Together, these findings support a model in which dysregulated host immune responses, rather than viral burden per se, are primary determinants of disease severity^33,34^.

Although TLR7 is canonically positioned upstream of IRF7 in endosomal RNA sensing pathways, our data do not support a dominant role for TLR7 in regulating the interferon responses downstream IRF7 during influenza infection. IRF7 expression and downstream antiviral programs remained largely intact in the absence of TLR7, suggesting compensation through alternative pathways, including type I interferon feedback loops and cytosolic RNA sensors such as RIG-I^24,35,36^(Supplementary Fig. 3). These findings demonstrate that transcriptional induction of a receptor does not necessarily imply functional dependency on downstream effects. Consistent with this, Jeisy-Scott *et al.* have also demonstrated that TLR7 recognition is dispensable for protective immunity during influenza infection, as *Tlr7^⁻/⁻^* mice are capable of mounting protective innate responses and controlling infection despite alterations in B cell responses^25^. Importantly, no naturally occurring mutations or variants in TLR7 have been reported to be associated with increased susceptibility to or severity of IAV in humans. In contrast, in SARS-CoV-2 infection, human genetic studies have demonstrated that TLR7 deficiency is strongly associated with severe COVID-19. Multiple studies in COVID-19 patients have identified rare deleterious TLR7 variants that impair type I interferon responses and predispose individuals to life-threatening disease highlighting its essential role in antiviral defense in that context^15–19^. These differences likely reflect distinct viral sensing requirements, with influenza relying more heavily on cytosolic pathways, whereas SARS-CoV-2 engages endosomal sensing mechanisms^23,35^.

In contrast to TLR7, IRF7 emerges as a critical and conserved regulator of innate immunity in both influenza and COVID-19. In our model, *Irf7* deficiency resulted in increased disease severity without affecting viral burden, suggesting that IRF7 promotes a response that protects the host and tissue integrity rather than direct viral clearance. Mechanistically, the interferons downstream of IRF7 not only restrict viral propagation but also preserve tissue integrity by constraining epithelial proliferation and coordinating repair responses^37^. The bronchial epithelial hyperplasia observed in *Irf7^−/−^* mice may reflect a compensatory yet pathological remodeling response to impaired interferon signaling, which requires further studies. Consistent with our findings, Wilk *et al*. demonstrated that deletion of *Irf7* renders the host highly susceptible to infection, associated with increased weight loss and reduced survival^38^. Similarly, Hatesuer *et al.* reported impaired type I interferon responses, increased weight loss, and reduced survival in *Irf7⁻/⁻* mice, highlighting its critical role in antiviral defense and immune regulation^39^. However, a recent study reported an improved outcome in *Irf7^⁻/⁻^* mice infected with a lethal dose of PR8^40^. This contradictory result is likely attributable to differences in viral dose, and experimental design. Notably, human genetic evidence further supports a central role for IRF7 in IAV infection, as a patient with a near-fatal infection caused by the 2009 pandemic H1N1 virus was found to carry two IRF7 loss-of-function mutated alleles impairing IRF7 protein function^41^. Another IAV infected patient with a rare E331V missense variant in IRF7 showed an impaired IRF7 activity, characterized by defective interferon priming and blunted antiviral responses in patient cells. Together, these findings establish IRF7 as a nonredundant determinant of antiviral host defense across respiratory viral infections.

Beyond innate immunity, our data also reveals distinct temporal roles for TLR7 and IRF7 in shaping adaptive immune responses. *Irf7^−/−^* mice exhibited a marked reduction in anti-HA IgG levels at early points (7 DPI), consistent with an impaired priming of adaptive immunity, as type I interferons downstream of IRF7 are known to enhance antigen presentation and T cell help^42,43^. This is further supported by prior studies demonstrating that the first direct stimulatory signal for local respiratory tract B cells during IAV infection is provided through the type I IFNR^44^. In contrast, *Tlr7^−/−^* mice showed a more pronounced defect at later stages (21 DPI), suggesting a role for TLR7 in the maturation and maintenance of the humoral response. TLR7 signaling in B cells has been implicated in germinal center formation, affinity maturation, and sustained antibody production^25,45^. Previous work demonstrated that although TLR7 signaling is dispensable for the generation of protective responses following primary IAV infection, it is required for optimal adaptive immune responses, including germinal center formation, antibody-secreting cell development, and long-term humoral memory, particularly in the context of vaccination^25^. Our findings provide evidence that IRF7 contributes to early adaptive immune priming, extending its role beyond innate antiviral defense. Together, these observations support functional division of labor in which IRF7 contributes to the linking between innate and adaptative immunity, whereas TLR7 enhances the optimization, quality, and durability of antibody response.

Several limitations of this study warrant consideration. The use of global knockout models precludes definitive assignment in specific cell populations for TLR7 and IRF7, and future studies employing conditional deletion strategies will be necessary to delineate their contributions to immune response within defined cellular compartments. Testing other influenza strains and respiratory viruses will help determine if our findings are broadly applicable. In summary, our study demonstrates that IRF7, rather than TLR7, is a critical determinant of disease outcome during influenza infection, functioning independently to regulate interferon responses, epithelial integrity, and early adaptive immune priming. While TLR7 is dispensable for acute antiviral control in IAV, it remains important for the development and maintenance of long-term adaptive immunity, a feature that appears to be shared across multiple viral contexts, including CoV2 infection^23^.

## Materials and Methods

### Mice

*Irf7^−/−^* (RBRC01420) mice were purchased from RIKEN, Japan. *Tlr7^−/−^*(008380) and C57BL/6 J (B6) were purchased from the Jackson Laboratory. All animals were housed in an animal facility at Tulane University.

### Study approval

All animal experiments were approved by the Institutional Animal Care and Use Committee at Tulane University. We have complied with all relevant ethical regulations for animal use.

### Influenza A infection

Mice were infected with H1N1 A/ PR/8/34 (PR8). Briefly, sublethal (50PFU) and lethal concentrations (83PFU) were diluted in 50ul of sterile saline, and mice were infected by oropharyngeal aspiration. Control, uninfected mice received sterile saline.

### Tissue collection and process

The health status of the mice was monitored daily. Euthanasia was performed if the mice exhibit a body weight loss of 25% or upon reaching the predetermined necropsy date. Blood was drawn via cardiac puncture into a BD microtainer and allowed to clot at room temperature for a minimum of 30 min. Subsequently, the blood samples were centrifuged at 3000 rpm for 10 min, and the serum was collected from the upper layer. One half of the lung was fixed in Z-FIX buffer at room temperature. The remaining lung tissue was divided as follows: half of the residual lung (left side) was immersed in 1 mL of Trizol reagent and stored at −80°C for RNA extraction; the other half of the residual lung (left side) was frozen on dry ice without any medium and stored at −80°C.

### Histological analysis and quantification of histopathologic lesions

Fixed tissues were processed, cut, and stained with hematoxylin and eosin (H&E) and digitally scanned with Axio Scan.Z1 (Zeiss, Thornwood, NY) by the Anatomic Pathology Core at Tulane National Biomedical Research Center. The histological injury level was assessed and scored by a pathologist.

### Immunohistochemistry

Tissue embedded in paraffin and fixed in formalin was sectioned at 4 μm, mounted on Superfrost Plus slides, and baked at 60°C for 3 h. Sections were deparaffinized in xylene, rehydrated through graded ethanol, and rinsed in distilled water. Heat-induced epitope retrieval was performed by boiling slides for 20 min in Tris-based buffer (pH 9; Vector Labs H-3301, 0.1% Tween-20), followed by transfer to citrate buffer (pH 6.0; Vector Labs H-3300) and cooling to room temperature. Slides were washed in PBS and loaded onto a Ventana Discovery Ultra autostainer for blocking, primary antibody incubation (rabbit anti-FluA, GeneTex GTX125989, 1:1000), secondary antibody incubation, and Rhodamine-based detection, followed by DAPI counterstaining. Slides were washed, mounted with Mowiol/DABCO aqueous medium, and imaged at 20× using a Zeiss Axio Scan.Z1 scanner.

### Digital image analysis

Slides were stained with DAPI and Influenza A-specific marker, as indicated before, is imaged across three channels, including an empty channel to account for autofluorescence. Influenza A-labeled cells were quantified using the HighPlex FL v4.2.14 module in HALO v3.6. Regions of interest were drawn around the entire lung section. Thresholds for positive detection were set by a veterinary pathologist based on fluorescent intensity in each channel and subsequently checked for accuracy by the same pathologist.

### RNA isolation

Tissues were collected in 1 mL of Trizol reagent (Invitrogen, Cat. No. 15596026) and RNA was extracted using the RNeasy Mini Kit (QIAGEN, Cat. No. 74104) according to the manufacturer’s protocol. The RNA concentration was measured using the NanoDrop 2000 spectrophotometer.

### Subgenomic M1 viral detection

Viral burden was determined by RT-PCR for viral matrix (M1) expression^46^.

### ELISA for Anti-HA detection

96-well plates were coated overnight at 4°C with recombinant HA protein (5 µg/mL; Influenza A/Puerto Rico/8/1934 H1N1 trimer, (Sino Biological Cat. No. 11684-V08H2) in PBS. Plates were washed with PBST and blocked for 2 h at room temperature with 1% BSA and 0.1% Tween-20 in PBS. Serum samples and controls were added in duplicate at predetermined dilutions, along with a standard curve of Influenza A H1N1 HA Monoclonal Antibody (Invitrogen, Cat. No. MA5-29929). After 1 h incubation and PBST washes, HRP-conjugated goat anti-mouse IgG (SouthernBiotech, Cat. No 1036-05) was added for 30 min at 37°C. Plates were washed, developed with TMB, stopped, and absorbance was read at 450 nm.

### Western Blot

Lung tissues were harvested from mice at the indicated time points (2 or 7 days post-infection), snap-frozen, and homogenized in RIPA Lysis and Extraction Buffer (Thermo Fisher Cat. No 89900), containing protease and phosphatase inhibitors (Cell Signaling, Cat. No 5871S). Protein concentration was determined using a BCA assay Buffer (Thermo Fisher Cat. No 23227). Equal amounts of protein (60 µg per lane) were resolved on 4 – 20% (BioRad, Cat. No 5671094) and transferred onto PVDF membranes (BioRad, Cat. No 1620177). Membranes were blocked in EveryBlot Blocking Buffer (BioRad, Cat. No 12010020). for 10 minutes at room temperature and incubated overnight with primary antibodies against IRF7 (Cell Signaling, Cat. No 72073), NF-κB (Cell Signaling, Cat. No 6956), IFN-α (Thermo Fisher Cat. No PA5-119649), IFN-γ (ProteinTech, Cat. No 29788-1-AP), RIG-I (Cell Signaling, Cat. No 4520), and β-actin (Cell Signaling, Cat. No 3700) After washing, membranes were incubated with HRP-conjugated secondary antibody (Cell Signaling, Cat. No 7074) for 1 h at room temperature. Protein bands were visualized using SuperSignal™ West Atto Ultimate Sensitivity Substrate (Thermo Fisher Cat. No A38556) in a ChemiDoc™ Imaging System (BioRad, Cat. No 12003153). Densitometric analysis was performed using Image J.

### Single-cell RNA sequencing

Lung was with forceps and small scissors and digested in 2 mL serum-free medium with 2 mg/mL collagenase (MilliporeSigma) and 80 U/mL DNase I (MilliporeSigma) for 60 min at 37 °C. 5000 live cells per sample were targeted by using 10× Single Cell RNAseq technology provided by 10× Genomics (10X Genomics). Full-length barcoded cDNAs were then generated and amplified by PCR to obtain sufficient mass for library construction. Pooled libraries at a final concentration of 1.8 pM were sequenced with paired-end single index configuration by Illumina NextSeq 550. Cell Ranger version 2.1.1 (10X Genomics) was used to process raw sequencing data and Loupe Cell Browser (10X Genomics) to obtain differentially expressed genes between specified cell clusters. In addition, Seurat suite version 2.2.1 was used for further quality control and downstream analysis. Filtering was performed to remove multiplets and broken cells, and uninteresting sources of variation were regressed. Variable genes were determined by iterative selection based on the dispersion versus average expression of the gene. For clustering, principal component (PC) analysis was performed for dimension reduction. The top 10 PCs will be selected by using a permutation-based test implemented in Seurat and passed to t-SNE for clustering visualization.

### Statistics and reproducibility

The results presented depict the output of independent mice (n = 1–5). Infected mice were analyzed at multiple time intervals post-infection to minimize bias and increase reproducibility. Data are shown as mean ± SEM. To compare values obtained from multiple groups over time, two-way analysis of variance (ANOVA) was used, followed by a Bonferroni post-hoc test. To compare values obtained from two groups, the unpaired Student’s *t*-test was performed. Statistical significance was taken at the P < 0.05 level.

## Notes

### Competing Interest Statement

The authors have declared no competing interest.

### Funding

- National Institutes of Health: P51OD011104-62 (X.Q.) and R01HL165265

## References

1 Currey, J. et al. Upregulation of inflammatory genes and pathways links obesity to severe COVID-19. Biochimica et Biophysica Acta (BBA) - Molecular Basis of Disease 1870, 167322 (2024).10.1016/j.bbadis.2024.167322

2 Álvarez, F., Froes, F., Rojas, A. G., Moreno-Perez, D. & Martinón-Torres, F. The challenges of influenza for public health. Future Microbiol 14, 1429–1436 (2019). 10.2217/fmb-2019-0203

3 Flerlage, T., Boyd, D. F., Meliopoulos, V., Thomas, P. G. & Schultz-Cherry, S. Influenza virus and SARS-CoV-2: pathogenesis and host responses in the respiratory tract. Nature Reviews Microbiology 19, 425–441 (2021). 10.1038/s41579-021-00542-7

4 Li, Y. et al. SARS-CoV-2 induces double-stranded RNA-mediated innate immune responses in respiratory epithelial-derived cells and cardiomyocytes. Proceedings of the National Academy of Sciences 118, e2022643118 (2021). doi:10.1073/pnas.2022643118

5 Datta, P. K., Liu, F., Fischer, T., Rappaport, J. & Qin, X. SARS-CoV-2 pandemic and research gaps: Understanding SARS-CoV-2 interaction with the ACE2 receptor and implications for therapy. Theranostics 10, 7448–7464 (2020). 10.7150/thno.48076

6 Bouvier, N. M. & Palese, P. The biology of influenza viruses. Vaccine 26 Suppl 4, D49–53 (2008). 10.1016/j.vaccine.2008.07.039

7 Sameer, A. S. & Nissar, S. Toll-Like Receptors (TLRs): Structure, Functions, Signaling, and Role of Their Polymorphisms in Colorectal Cancer Susceptibility. Biomed Res Int 2021, 1157023 (2021). 10.1155/2021/1157023

8 Diamond, M. S. & Kanneganti, T. D. Innate immunity: the first line of defense against SARS-CoV-2. Nat Immunol 23, 165–176 (2022). 10.1038/s41590-021-01091-0

9 Akira, S., Takeda, K. & Kaisho, T. Toll-like receptors: critical proteins linking innate and acquired immunity. Nature Immunology 2, 675–680 (2001). 10.1038/90609

10 Yang, M.-Y. et al. Role of toll-like receptors in the pathogenesis of COVID-19: Current and future perspectives. Scandinavian Journal of Immunology 98, e13275 (2023). 10.1111/sji.13275

11 Duan, T., Du, Y., Xing, C., Wang, H. Y. & Wang, R. F. Toll-Like Receptor Signaling and Its Role in Cell-Mediated Immunity. Front Immunol 13, 812774 (2022). 10.3389/fimmu.2022.812774

12 Oh, J. Z., Kurche, J. S., Burchill, M. A. & Kedl, R. M. TLR7 enables cross-presentation by multiple dendritic cell subsets through a type I IFN-dependent pathway. Blood 118, 3028–3038 (2011). 10.1182/blood-2011-04-348839

13 Zhang, S.-Y. et al. TLR3 immunity to infection in mice and humans. Current Opinion in Immunology 25, 19–33 (2013). 10.1016/j.coi.2012.11.001

14 Spiering, A. E. & de Vries, T. J. Frontiers | Why Females Do Better: The X Chromosomal TLR7 Gene-Dose Effect in COVID-19. Frontiers in Immunology 12 (2021/11/11). 10.3389/fimmu.2021.756262

15 X, S., et al. Genetic Screening for TLR7 Variants in Young and Previously Healthy Men With Severe COVID-19 - PubMed. Frontiers in immunology 12 (07/23/2021). 10.3389/fimmu.2021.719115

16 CI, v. d. M., et al. Presence of Genetic Variants Among Young Men With Severe COVID-19 - PubMed. JAMA 324 (08/18/2020). 10.1001/jama.2020.13719

17 Naushad, S. M. et al. The role of TLR7 agonists in modulating COVID-19 severity in subjects with loss-of-function TLR7 variants. Scientific Reports 2023 13:1 13 (2023-08-11). 10.1038/s41598-023-40114-8

18 T, A., et al. X-linked recessive TLR7 deficiency in ∼1% of men under 60 years old with life-threatening COVID-19 - PubMed. Science immunology 6 (08/19/2021). 10.1126/sciimmunol.abl4348

19 C, F., et al. Association of Toll-like receptor 7 variants with life-threatening COVID-19 disease in males: findings from a nested case-control study - PubMed. eLife 10 (03/02/2021). 10.7554/eLife.67569

20 Ma, W., Huang, G., Wang, Z., Wang, L. & Gao, Q. Frontiers | IRF7: role and regulation in immunity and autoimmunity. Frontiers in Immunology 14 (2023/08/10). 10.3389/fimmu.2023.1236923

21 TM, C., et al. Respiratory viral infections in otherwise healthy humans with inherited IRF7 deficiency - PubMed. The Journal of experimental medicine 219 (07/04/2022). 10.1084/jem.20220202

22 Strunz, B. et al. Type I Interferon Autoantibodies Correlate With Cellular Immune Alterations in Severe COVID-19. The Journal of Infectious Diseases 230 (2024/08/16). 10.1093/infdis/jiae036

23 C, W., et al. Deficiency of Tlr7 and Irf7 in mice increases the severity of COVID-19 through the reduced interferon production - PubMed. Communications biology 7 (09/17/2024). 10.1038/s42003-024-06872-5

24 Pang, I. K., Pillai, P. S. & Iwasaki, A. Efficient influenza A virus replication in the respiratory tract requires signals from TLR7 and RIG-I. Proc Natl Acad Sci U S A 110, 13910–13915 (2013). 10.1073/pnas.1303275110

25 Jeisy-Scott, V. et al. TLR7 Recognition Is Dispensable for Influenza Virus A Infection but Important for the Induction of Hemagglutinin-Specific Antibodies in Response to the 2009 Pandemic Split Vaccine in Mice. Journal of Virology 86 (2012-10-15). 10.1128/jvi.01064-12

26 Rappe, J. C. F. et al. A TLR7 antagonist restricts interferon-dependent and - independent immunopathology in a mouse model of severe influenza. J Exp Med 218 (2021). 10.1084/jem.20201631

27 Miles, M. A. et al. TLR7 Promotes Acute Inflammatory-Driven Lung Dysfunction in Influenza-Infected Mice but Prevents Late Airway Hyperresponsiveness. International Journal of Molecular Sciences 25, 13699 (2024).

28 Yin, Q. et al. A TLR7-nanoparticle adjuvant promotes a broad immune response against heterologous strains of influenza and SARS-CoV-2. Nature Materials 22, 380–390 (2023). 10.1038/s41563-022-01464-2

29 Ciancanelli, M. J. et al. Infectious disease. Life-threatening influenza and impaired interferon amplification in human IRF7 deficiency. Science 348, 448–453 (2015). 10.1126/science.aaa1578

30 Wang, C. et al. COVID-19 and influenza infections mediate distinct pulmonary cellular and transcriptomic changes. Communications Biology 2023 6:1 6 (2023-12-13). 10.1038/s42003-023-05626-z

31 Tabula Muris, C., et al. Single-cell transcriptomics of 20 mouse organs creates a Tabula Muris. Nature 562, 367–372 (2018). 10.1038/s41586-018-0590-4

32 Seo, S.-U. et al. MyD88 Signaling Is Indispensable for Primary Influenza A Virus Infection but Dispensable for Secondary Infection. Journal of Virology 84, 12713–12722 (2010). 10.1128/jvi.01675-10

33 W, W., et al. IRF7 Is Required for the Second Phase Interferon Induction during Influenza Virus Infection in Human Lung Epithelia - PubMed. Viruses 12 (03/29/2020). 10.3390/v12040377

34 Fan, S., Popli, S., Chakravarty, S., Chakravarti, R. & Chattopadhyay, S. Non-transcriptional IRF7 interacts with NF-κB to inhibit viral inflammation. Journal of Biological Chemistry 300 (2024/04/01). 10.1016/j.jbc.2024.107200

35 Coch, C. et al. RIG-I Activation Protects and Rescues from Lethal Influenza Virus Infection and Bacterial Superinfection. Molecular Therapy 25 (2017 Jul 8). 10.1016/j.ymthe.2017.07.003

36 Kandasamy, M. et al. RIG-I Signaling Is Critical for Efficient Polyfunctional T Cell Responses during Influenza Virus Infection. PLOS Pathogens 12 (Jul 20, 2016). 10.1371/journal.ppat.1005754

37 Major, J. et al. Type I and III interferons disrupt lung epithelial repair during recovery from viral infection. Science 369, 712–717 (2020). 10.1126/science.abc2061

38 Wilk, E. et al. RNAseq expression analysis of resistant and susceptible mice after influenza A virus infection identifies novel genes associated with virus replication and important for host resistance to infection. BMC Genomics 16 (2015). 10.1186/s12864-015-1867-8

39 B, H., et al. Deletion of Irf3 and Irf7 Genes in Mice Results in Altered Interferon Pathway Activation and Granulocyte-Dominated Inflammatory Responses to Influenza A Infection - PubMed. Journal of innate immunity 9 (2017). 10.1159/000450705

40 Cui, J. et al. *Irf7* Deficiency Confers Protection Against Influenza Infection, Independent of irf3. International Journal of Biological Sciences 22, 1974–1996 (2026). 10.7150/ijbs.126714

41 Ciancanelli, M. J. et al. Life-threatening influenza and impaired interferon amplification in human IRF7 deficiency. Science 348, 448–453 (2015). 10.1126/science.aaa1578

42 Le Bon, A. et al. Type I Interferons Potently Enhance Humoral Immunity and Can Promote Isotype Switching by Stimulating Dendritic Cells In Vivo. Immunity 14, 461–470 (2001). 10.1016/s1074-7613(01)00126-1

43 Gessani, S., Conti, L., Del Cornò, M. & Belardelli, F. Type I Interferons as Regulators of Human Antigen Presenting Cell Functions. Toxins 6, 1696–1723 (2014). 10.3390/toxins6061696

44 ES, C., WL, C. & N, B. Type I IFN receptor signals directly stimulate local B cells early following influenza virus infection - PubMed. Journal of immunology (Baltimore, Md.: 1950) 176 (04/01/2006). 10.4049/jimmunol.176.7.4343

45 C, S., et al. B cell-intrinsic TLR7 signaling is essential for the development of spontaneous germinal centers - PubMed. Journal of immunology (Baltimore, Md.: 1950) 193 (11/01/2014). 10.4049/jimmunol.1401720

46 Crowe, C. R. et al. Critical Role of IL-17RA in Immunopathology of Influenza Infection. The Journal of Immunology 183 (2009/10/15). 10.4049/jimmunol.0900995

