## Supplemental Figures and Tables for "IRF7 deficiency increases disease severity independently of TLR7 recognition in Influenza A infection in mice"

**A.**

| Tissue | Mean | Median | Median > 0 | # cells | % 0 | % > 0 |
|---|---|---|---|---|---|---|
| non-classical monocyte | 0.42 | 0 | 1.15 | 220 | 65.00 | 35.00 |
| classical monocyte | 0.27 | 0 | 0.77 | 161 | 68.94 | 31.06 |
| alveolar macrophage | 0.20 | 0 | 0.92 | 345 | 78.26 | 21.74 |
| myeloid cell | 0.06 | 0 | 0.79 | 87 | 91.95 | 8.05 |
| B cell | 0.04 | 0 | 1.34 | 204 | 97.06 | 2.94 |
| leukocyte | 0.03 | 0 | 1.09 | 152 | 97.37 | 2.63 |
| T cell | 0.03 | 0 | 1.00 | 247 | 97.57 | 2.43 |
| lung endothelial cell | 0.01 | 0 | 0.84 | 462 | 99.35 | 0.65 |
| NA | 0.00 | 0 | NA | 45 | 100.00 | 0.00 |
| stromal cell | 0.00 | 0 | 1.05 | 2540 | 99.84 | 0.16 |
| type II pneumocyte | 0.00 | 0 | NA | 89 | 100.00 | 0.00 |
| natural killer cell | 0.00 | 0 | NA | 824 | 100.00 | 0.00 |
| mast cell | 0.00 | 0 | NA | 24 | 100.00 | 0.00 |
| ciliated columnar cell of tracheobronchial tree | 0.00 | 0 | NA | 49 | 100.00 | 0.00 |

**B**

| Tissue | Mean | Median | Median > 0 | # cells | % 0 | % > 0 |
|---|---|---|---|---|---|---|
| myeloid cell | 0.12 | 0 | 1.10 | 87 | 87.36 | 12.64 |
| lung endothelial cell | 0.12 | 0 | 1.16 | 462 | 90.04 | 9.96 |
| alveolar macrophage | 0.06 | 0 | 0.86 | 345 | 92.75 | 7.25 |
| stromal cell | 0.04 | 0 | 1.19 | 2540 | 96.46 | 3.54 |
| ciliated columnar cell of tracheobronchial tree | 0.03 | 0 | 1.39 | 49 | 97.96 | 2.04 |
| type II pneumocyte | 0.03 | 0 | 0.68 | 89 | 96.63 | 3.37 |
| classical monocyte | 0.01 | 0 | 0.69 | 161 | 98.14 | 1.86 |
| B cell | 0.01 | 0 | 1.61 | 204 | 99.51 | 0.49 |
| leukocyte | 0.01 | 0 | 0.94 | 152 | 99.34 | 0.66 |
| NA | 0.00 | 0 | NA | 45 | 100.00 | 0.00 |
| non-classical monocyte | 0.00 | 0 | 1.06 | 220 | 99.55 | 0.45 |
| natural killer cell | 0.00 | 0 | 0.78 | 824 | 99.76 | 0.24 |
| T cell | 0.00 | 0 | NA | 247 | 100.00 | 0.00 |
| mast cell | 0.00 | 0 | NA | 24 | 100.00 | 0.00 |

**Supplemental Table S1. Summary statistics of *Tlr7* and *Tlr3* expression across lung cell populations. (A)** Quantitative summary of *Tlr7* expression across annotated lung cell populations from the droplet-based single-cell RNA-seq dataset of the Tabula Muris Consortium[31]. For each cell type, mean expression, median expression, median expression among expressing cells (>0), total number of cells analyzed, percentage of cells with zero expression (% 0), and percentage of cells with detectable expression (% >0) are shown. **(B)** Quantitative summary of *Tlr3* expression across the same lung cell populations. Metrics are defined as in (A).

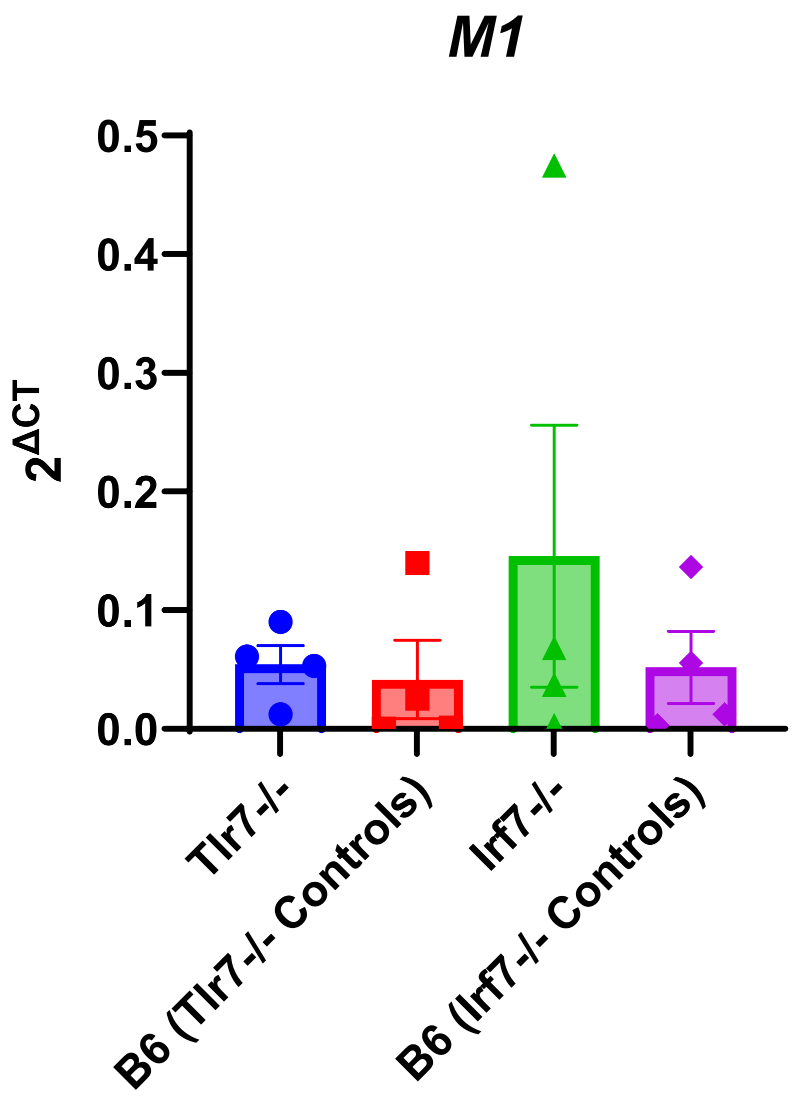


**Supplemental Figure 1. M1 viral gene expression is not altered by Tlr7 or Irf7 deficiency.** Relative expression of influenza A virus matrix gene (*M1*) in lung tissue from wild-type (B6), *Tlr7* knockout mice, and *Irf7* knockout mice. Viral RNA levels were quantified by qPCR and normalized using the 2^-ΔCt method. Each point represents an individual mouse; bars indicate mean ± SEM. No significant differences in *M1* expression were observed between groups, indicating comparable viral burden across genotypes.

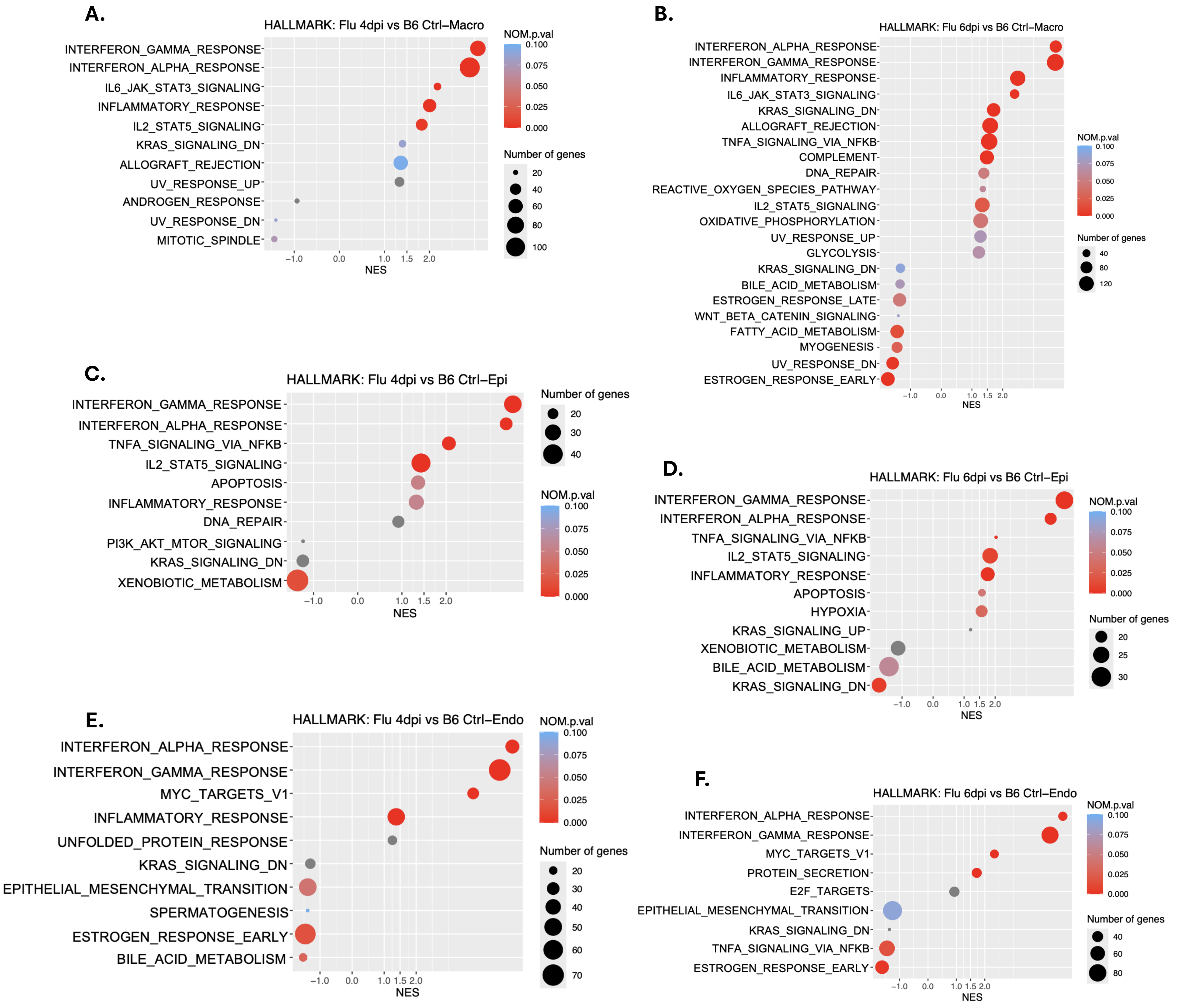


**Supplemental Figure 2. GSEA. A–B)** Gene set enrichment analysis (GSEA) of Hallmark pathways in macrophages from IAV infected versus B6 control mice at 4 dpi (A) and 6 dpi (B). **C–D)** GSEA of Hallmark pathways in epithelial cells comparing IAV infected and B6 control mice at 4 dpi (C) and 6 dpi (D). **E–F)** GSEA of Hallmark pathways in endothelial cells comparing IAV infected and B6 control mice at 4 dpi (E) and 6 dpi (F). Normalized enrichment scores (NES) are shown on the x-axis. Dot size represents the number of genes contributing to each pathway, and color indicates normalized *p*-value (NOM p.val)

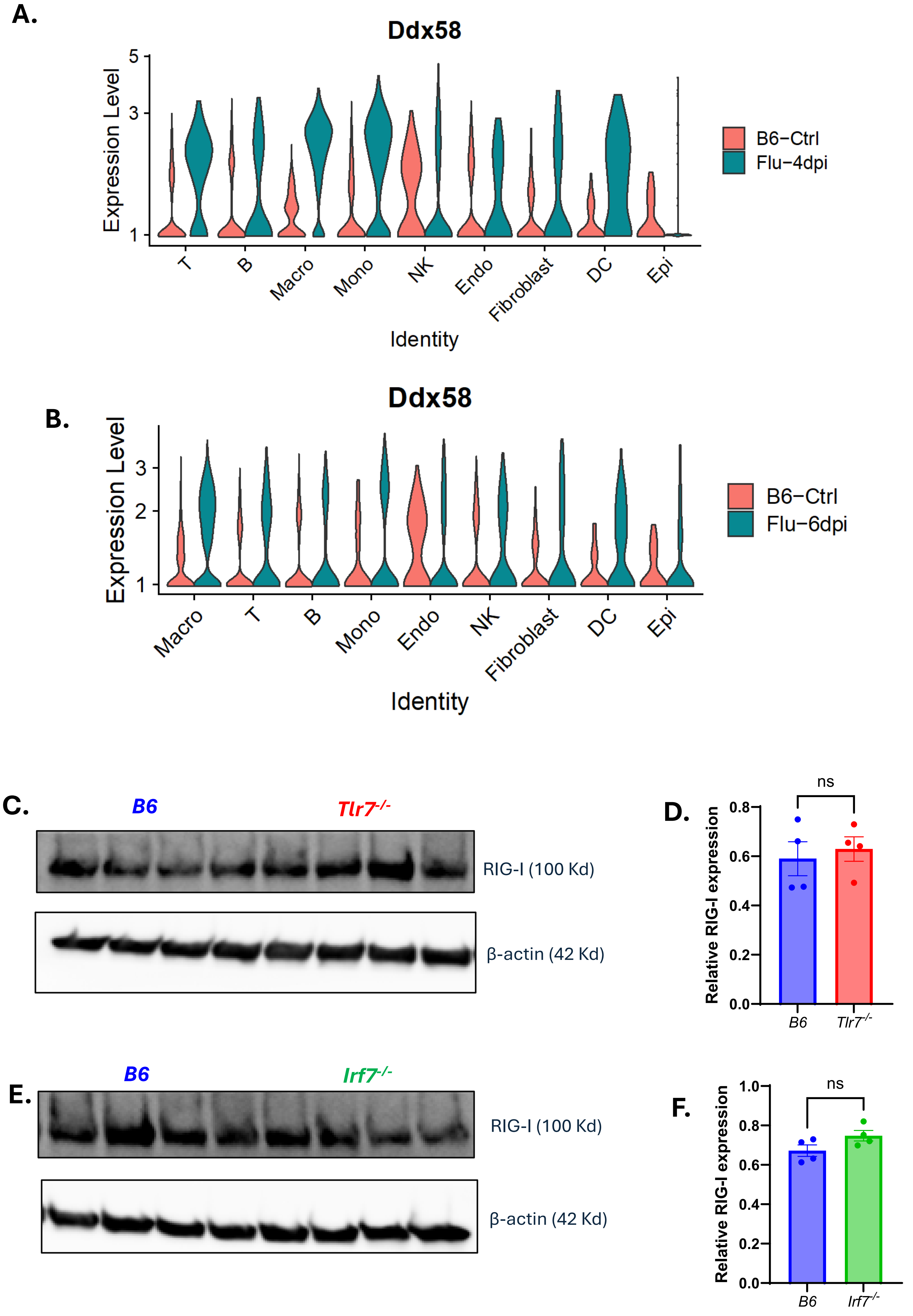


**Supplemental Figure 3. RIG-I expression in IAV infection. A - B)** Single-cell RNA-seq analysis showing Ddx58 (RIG-I) expression across indicated lung cell populations in B6 control and influenza A infected mice at 4 dpi (A) and 6 dpi (B). **C–F)** Western blots and corresponding densitometric quantification of RIG-I (100 kDa) in lung lysates from B6 and *Tlr7*$^{-/-}$ (C–D) or *Irf7*$^{-/-}$ (E–F) mice. β-actin (42 kDa) is shown as a loading control. Protein levels were normalized to β-actin and expressed relative to B6 controls. Data are presented as mean ± SEM. Statistical significance was determined by unpaired two-tailed Student's *t*-test; ns, *not significant*.
